# Evidence for anti-viral effects of complete Freund’s adjuvant in the mouse model of enterovirus infection

**DOI:** 10.1101/2020.05.27.120121

**Authors:** Arunakumar Gangaplara, Chandirasegaran Massilamany, Ninaad Lasrado, David Steffen, Jay Reddy

**Author notes:** Laboratory of Early Sickle Mortality Prevention, Cellular and Molecular Therapeutics Branch, National Heart, Lung, and Blood Institute, National Institutes of Health, Bethesda, Maryland, 20892, United States of America; (A.G). CRISPR Therapeutics, Cambridge, Massachusetts, 02139, United States of America; (C.M). Equal contributors (Chandirasegaran Massilamany, Ninaad Lasrado).

## Abstract

Group B Coxsackieviruses belonging to the genus, Enterovirus, contain six serotypes that induce various diseases, whose occurrence may involve the mediation of more than one serotype. We recently identified immunogenic epitopes within CVB3 viral protein 1 that induce anti-viral T cell responses in mouse models of CVB infections. In our investigations to determine the protective responses of the viral epitopes, we unexpectedly noted that animals immunized with complete Freund’s adjuvant (CFA) alone and later challenged with CVB3 were completely protected against myocarditis. Similarly, the pancreatitis-inducing ability of CVB3 was remarkably reduced to only 10% in the CFA group as opposed to 73.3% in the control group that received no CFA. Additionally, no mortalities were noted in the CFA group, whereas 40% of control animals died during the course of 21 days post-infection with CVB3. Taken together, our data suggest that the adjuvant effects of CFA may be sufficient for protection against CVB infections. These observations may provide new insights into our understanding of the occurrence of viral infections. One example is Coronavirus disease-19 (COVID-19) as individuals suffering from COVID-19 who have been vaccinated with Bacillus Calmette–Guérin appear to have fewer morbidities and mortalities than unvaccinated individuals.

## Introduction

Enteroviruses belonging to the *Picornaviridae* family are positive-sense, single-stranded RNA viruses. Based on the currently adopted method of molecular typing of the viral protein (VP)1 nucleotide composition, 13 species of enteroviruses have been identified [1]. Infections caused by four enterovirus species, Enterovirus A to D, are the most common that occur in humans, especially infants (children less than 1 year of age) and immune-compromised individuals [1, 2]. Enteroviruses induce a wide spectrum of illnesses, such as meningitis, encephalitis, paralysis, myocarditis, and rash/foot and mouth disease [2]. Although enteroviral infections can occur anywhere in the world, recent outbreaks of respiratory illness in the United States highlight their growing importance in human health [3–5].

We have been studying the cellular and molecular mechanisms of protective immune responses in mouse models, particularly for Coxsackievirus B3 (CVB3) and CVB4, which are implicated in the causation of myocarditis/dilated cardiomyopathy and Type I diabetes (TID), respectively [6, 7]. In our efforts to determine the protective effects of viral epitopes, we unexpectedly noted that animals immunized with complete Freund’s adjuvant (CFA) containing *Mycobacterium tuberculosis* (M. tb) extract were found to be completely protected from CVB3 infection. Our data may provide new insights regarding the occurrence of viral infections such as Coronavirus disease-19 (COVID-19), since individuals vaccinated with Bacillus Calmette–Guérin (BCG) tend to show fewer morbidities and mortalities than unvaccinated individuals [8, 9].

## Materials and Methods

### Mice

Six-to-eight-week old, female A/J mice (H-2^a^) were procured from the Jackson Laboratory (Bar Harbor, ME, USA). Animals were maintained according to the institutional guidelines of the University of Nebraska-Lincoln (UNL), Lincoln, NE, and approval for animal studies was granted by the Institutional Animal Care and Use Committee, UNL (protocol #1904, approved January 2, 2020). Mice infected with CVB3 were monitored closely for clinical signs suggestive of distress. All research staff followed biosafety level 2 guidelines while handling the animals. Animals whose clinical signs persisted, did not eat or drink, and failed to move when touched or prodded physically were immediately euthanized. Euthanasia was performed using a carbon dioxide chamber as recommended by the Panel on Euthanasia, the American Veterinary Medical Association.

### Virus propagation and infection

The Nancy strain of CVB3 was procured from the American Type Culture Collection (ATCC, Manassas, VA, USA), and the virus was titrated in Vero cells (ATCC). The adherent Vero cells were grown to 80 to 90% confluence in 75cm^2^ flasks in EMEM/10% fetal bovine serum (FBS) and were later infected with CVB3 with multiplicity of infection 1 in EMEM containing no FBS. After incubation at 37° C for 1 hour with gentle intermittent rocking, maintenance medium (EMEM/2% FBS) was added. Based on the cytopathic effect of virus during the next 1 to 2 days, supernatants containing virus were harvested. After determining 50% tissue culture infective dose (TCID50) values based on the Reed-Muench method, the virus stocks were aliquoted and preserved at −80° C [10]. To infect mice, virus stock diluted in 1x PBS to contain 50 TCID50 in 100 μl was administered intraperitoneally (i.p.). We chose this dose based on titration experiments [10] that allowed us to capture pathological changes in both heart and pancreas over a period of 3 weeks by avoiding acute mortalities that usually occur at relatively higher doses within ~10 days post-infection [11, 12]. Animals were monitored closely, cages were changed once in 2 days, and body weights were taken daily until termination. In addition, an alternative food and fluid source, trans gel diet (ClearH2O, Portland, ME, USA), was placed on the cage floor as needed.

### Challenge studies in animals immunized with CFA

Our initial focus was to determine the protective effects, if any, of virus-reactive T cells by immunizing mice with viral peptides that we have recently described elsewhere [13]. In these investigations, two groups – CVB3 infection alone and CFA immunizations/CVB3 challenge – were involved. Groups of mice were immunized subcutaneously with or without CFA containing M. tb [14] (Difco Laboratories, Detroit, MI, USA) to a final concentration of 5 mg/ml as a single dose (200 μl; Sigma-Aldrich, St. Louis, MO, USA) on day −7 in sternal and inguinal regions [15]. Seven days later (day 0), animals were challenged with CVB3 at 50 TCID50/mouse, i.p., and after taking body weights and monitoring for mortalities, experiments were terminated on days 20-21 post-infection and tissues were collected for histology.

### Histology

Hearts and pancreata were fixed in 10% phosphate-buffered formalin and processed to obtain 5 μm thick serial sections, ~50 μm apart. All sections were stained by Hematoxylin and Eosin (H & E). The analysis was performed by a board-certified pathologist blinded to treatment, and total number of inflammatory foci were obtained as reported previously [10, 16].

### Statistics

Generalized linear mixed models were used to analyze the data pertaining to body weights and survival curves using Proc Glimmix in SAS (Version 9.3, SAS Institute Inc., Cary, NC, USA). Graphs were prepared by GraphPad Prism software Version 8.0 (GraphPad Software, Inc. La Jolla, CA, USA). Barnard’s exact test was used to analyze the histological parameters [17].

## Results and Discussion

In this report, we provide evidence that immunization with immune-stimulating adjuvants like CFA alone can offer protection against viral infections. In this setting, we used A/J mice that are highly susceptible to CVB infections, with affected animals showing severe pancreatitis and myocarditis within approximately 7 to 10 days post-infection [13, 16, 18]. Although our primary focus was to determine whether viral peptides can offer protection in challenge studies with CVB3, we made an unexpected observation that the protection offered by CFA emulsions containing viral peptides was indistinguishable (data not shown) from that conferred by immunization with CFA alone. First, we noted that the positive control group (unimmunized) infected with CVB3 showed reduction in body weights by ~20% as expected, whereas none of the animals immunized with CFA alone lost body weight (Fig 1a). Second, mortality patterns were also found to be similar to those of body weights, in that none of the animals immunized with CFA alone died, whereas 40% of animals (6/15) in the control group died (Fig 1b), pointing to the possibility that CFA-immunized animals would also be free of histologic disease.

**Figure 1.**
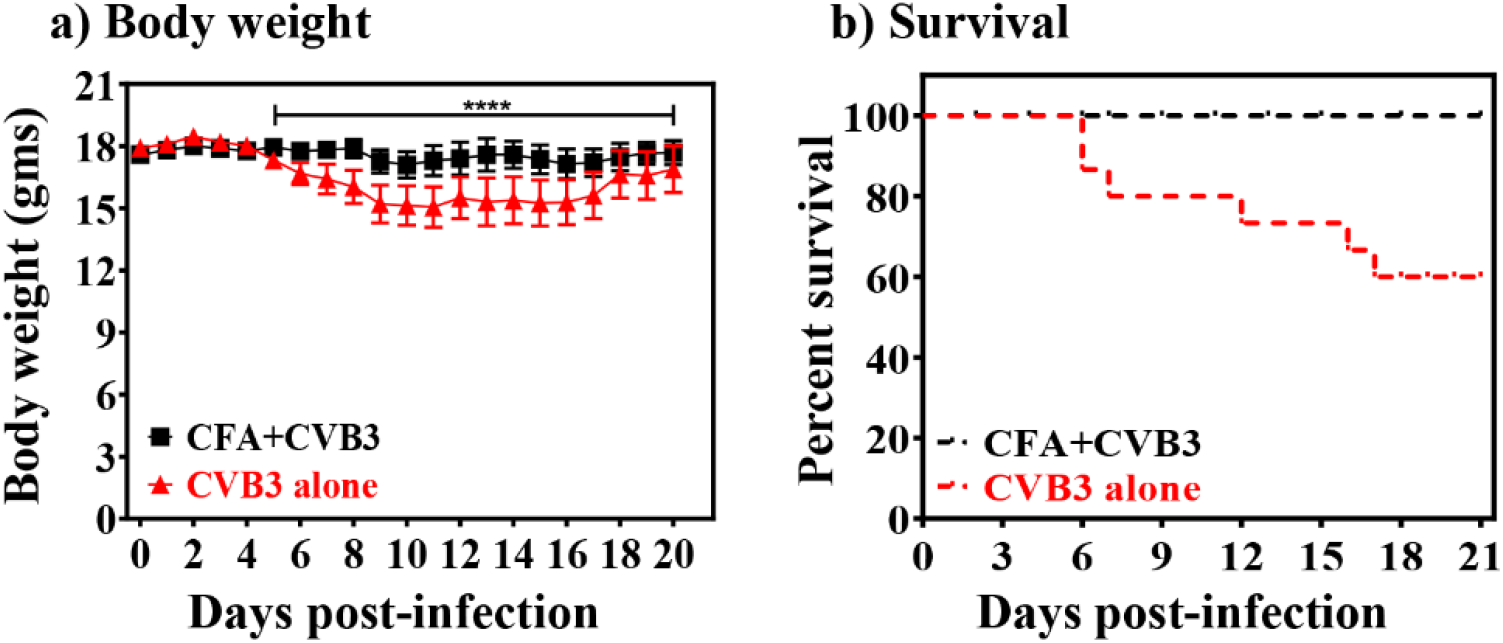
Evaluation of effects of CFA in CVB3 infection of A/J mice. **a) Body weight**. A/J mice were immunized once (-D7) with or without CFA alone and were challenged with CVB3 on day 0. Body weights were taken up to 20 days post-challenge with CVB3. Mean ± SEM values obtained from two experiments, each involving 5 to 10 mice, are shown. **b) Survival.** Mortalities noted up to 21 days post-challenge are shown in the survival curves. ****p≤0.0001.

To investigate pathological changes, we examined the hearts and pancreata by H & E staining and scored disease severity as we have described previously [10, 15, 16]. As indicated in Table 1, top panel, and Figure 2, heart sections in 40% (6/15) of the animals from the CVB3-infected group showed myocardial lesions containing inflammatory foci with macrophage infiltrates, necrosis and mineralization as expected [10, 16], but none of the animals in the CFA group had any detectable lesions. Similarly, histological evaluation of pancreatic sections from the CVB3-infected group revealed expected lesions such as atrophy, inflammation, necrosis, and mineralization, whereas in the CFA group, only 10% (1/10) of the animals had detectable lesions (Table 1, bottom panel, and Figure 2). Taken together, the findings that animals immunized with CFA alone were protected against both myocarditis and pancreatitis induced with CVB3 supports the idea that non-specific priming of the immune system with adjuvants like CFA may be sufficient to prevent virus infections.

**Figure 2.**
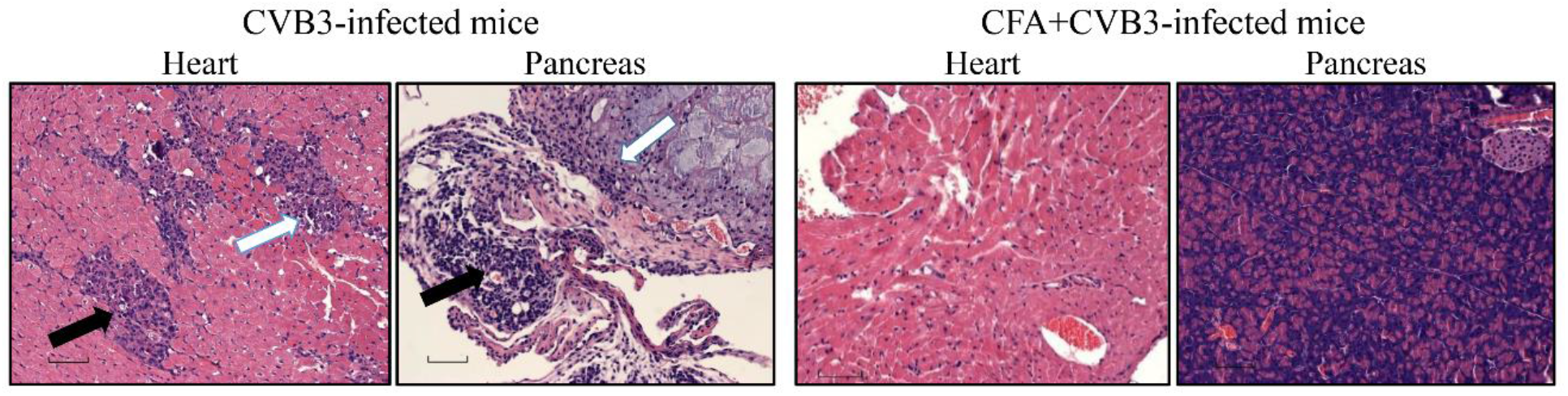
Determination of histological changes in mice immunized with or without CFA and later challenged with CVB3. A/J mice were immunized with or without CFA on D-7 and animals were challenged with CVB3 i.p., on day 0. Hearts and pancreata were examined by H and E staining to determine histological changes. The left panel indicates representative sections from CVB3-infected mice showing multiple inflammatory foci, and necrosis (solid arrow) and mineralization (empty arrow) in the heart, whereas pancreas showed changes such as atrophy and infiltrations (solid arrow) and necrosis and mineralization (empty arrow). The right panel denotes heart and pancreatic sections from CFA+CVB3-infected group in which lesions were absent. Original magnification: 20x.

**Table 1:**
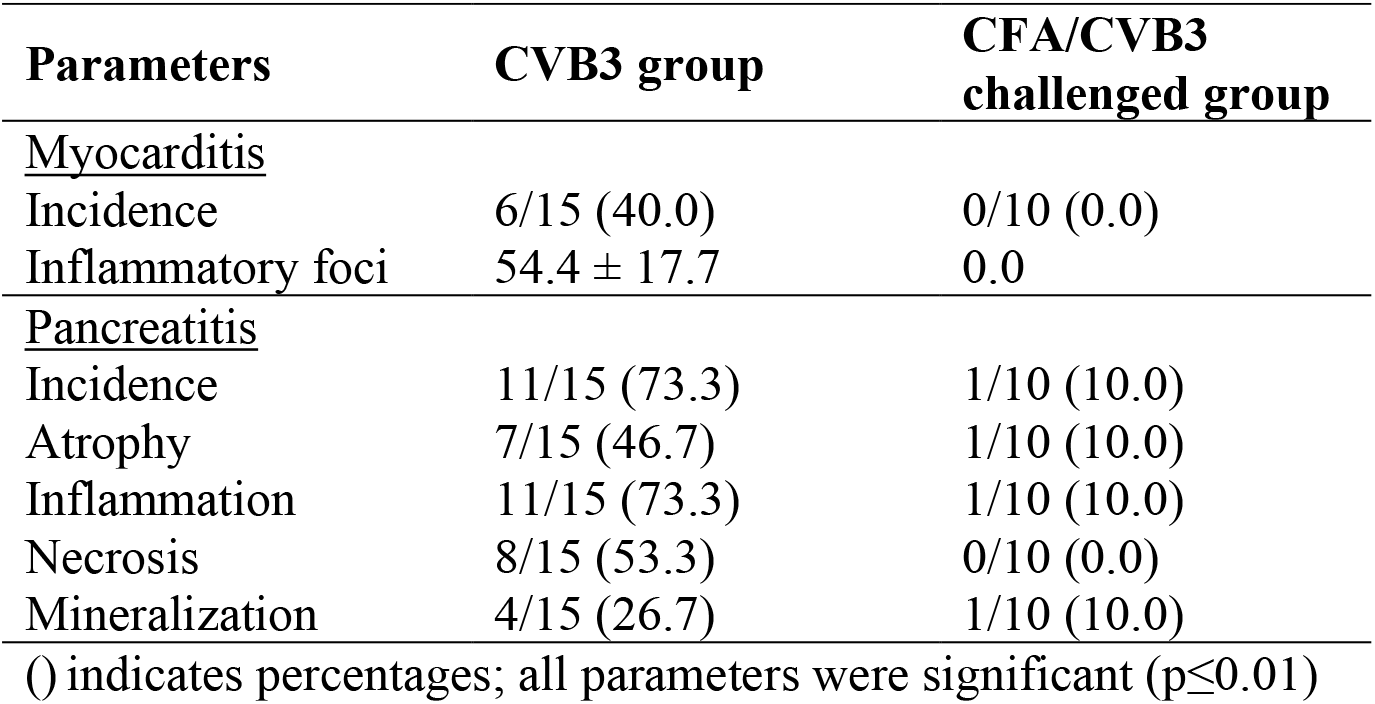
Histological evaluation of hearts and pancreata in mice immunized with or without CFA challenged with CVB3.

| Parameters | CVB3 group | CFA/CVB3 challenged group |
| --- | --- | --- |
| <u>Myocarditis</u> |  |  |
| Incidence | 6/15 (40.0) | 0/10 (0.0) |
| Inflammatory foci | 54.4 ± 17.7 | 0.0 |
| <u>Pancreatitis</u> |  |  |
| Incidence | 11/15 (73.3) | 1/10 (10.0) |
| Atrophy | 7/15 (46.7) | 1/10 (10.0) |
| Inflammation | 11/15 (73.3) | 1/10 (10.0) |
| Necrosis | 8/15 (53.3) | 0/10 (0.0) |
| Mineralization | 4/15 (26.7) | 1/10 (10.0) |

() indicates percentages; all parameters were significant ( $p \leq 0.01$ )

Disease protection offered by CFA was rather perplexing to explain, because CFA was not expected to induce antigen-specific immune responses to prevent infection with CVB3. Historically, CFA containing M. tb has been used as a powerful immune-stimulating adjuvant, and it contains immunoreactive molecules, such as N-acetylmuramyl-l-alanyl-d-isoglutamine (muramyl dipeptide) and trehalose 6,6’-dimycolate, all of which promote Th1 cell polarization by inducing IFN-γ [19–21]. Reports indicate that the BCG bacterium can enhance immunogenicity and also promote type 1 IFN response as shown in various settings such as vaccinations against influenza and hepatitis B viruses in humans [22, 23], and infection studies with encephalomyocarditis, murine hepatitis (mouse coronavirus), type 1 and 2 herpes simplex, vaccinia and foot-and-mouth disease viruses in mice [24–29]. Additionally, bacterial CpG nucleotides promote Th1 (IFN-γ) response [30], which is critical for protection against intracellular pathogens and viruses through the production of interleukin-12 by interacting with toll-like receptor-9 [31, 32]. Thus, we believe that the disease-protective ability of CFA may reflect non-antigen-specific effects attributable to the adjuvanticity of M. tb.

However, our data point to a few other possible mechanisms: (i) Exposure to non-specific infections that promote Th1 cytokine responses may offer protection against viral infections possibly by preventing viral replication [20, 21, 33]. If this holds true, then our data may also potentially provide credence to the hygiene hypothesis [34, 35]. (ii) A growing body of evidence suggests that the trained innate memory may be an important property of the innate immune system. Myeloid cells, such as monocytes and macrophages, Natural killer (NK) cells, NK-T cells, γδ T cells, and possibly innate lymphoid cells, exposed to a microbe – for example, ‘x’ microbe – can robustly respond to this microbe upon re-exposure, and also for other unrelated microbial stimulations through epigenetic and metabolic reprogramming pathways [36–38]. Consistent with this notion, it has been shown that BCG vaccine can offer protection against other unrelated pathogens, such as *Candida albicans, Schistosoma mansoni* and *Staphylococcus aureus* [39–42], including autoimmune diseases such as Type I Diabetes and multiple sclerosis [43–47]. Conversely, it is also possible that exposure to one type of pathogen can suppress immune responses to entirely different types of pathogens. For example, co-administration of oral polio vaccine with BCG at birth can diminish the response to purified protein derivative from BCG [48]. Although whether or not it is currently known that trained innate memory is an underlying mechanism for CFA effects as noted in our studies, this aspect may need to be investigated.

Finally, it may be noted that the ongoing pandemic outbreak of COVID-19 has offered new insights in regard to BCG vaccination, in that morbidities and mortalities appear to be low in BCG vaccine recipients [8]. Although the practice of BCG vaccination varies widely across the globe [49, 50], isolated reports showing an evidence of an inverse relationship between BCG vaccinations and COVID-19 attributable mortalities as noted in few countries, if not all (Supplementary Table 1) [8, 9]. But, such a relationship could be contributed by the confounding factors such as demographic and genetic variations in the affected populations, loopholes in the social distancing and quarantine measures, availability of diagnostic tools and prompt reporting of positive cases [51]. Nonetheless, the evolving notion of inverse relationship between BCG and clinical outcomes of severe acute respiratory syndrome coronavirus 2 (SARS CoV-2) infection that causes COVID-19 raise fundamental questions as to the immunological relationship between the two entities. While the BCG vaccine contains live *Mycobacterium bovis* (a pathogen of cattle), CFA contains the killed extract of M. tb (a pathogen of humans). Yet all 13 known mycobacterial species, including the two species identified above, show more than 99% nucleotide similarity, suggesting that all of them may have similar adjuvant properties [52, 53]. Thus, our data may provide experimental evidence for the notion that certain degree of resistance to COVID-19 infection in BCG vaccine recipients may be attributable to the adjuvant effects of mycobacteria that may involve the trained innate memory. If this notion holds true, then it creates opportunities to use BCG or its ingredients, such as CpG nucleotides, as anti-viral compounds. To this end, several phase III clinical trials have been initiated in a number of countries to test whether the BCG vaccination can offer protection or alter prognosis of COVID-19 infection [51, 54]. However, it should be noted that BCG vaccination is performed at birth, whereas COVID-19 infection tends to be more common in the elderly population [55, 56]. Although existence of multiple co-morbidities may explain fatalities in these patient populations, the data also lead to the question whether beneficial effects of BCG can last so long. Additionally, readers are urged to cautiously interpret the direct translational significance of our findings to humans, since CVB3 and SARS CoV-2 are two different viruses. Thus, evaluation of the effect of BCG in appropriate animal models that capture the disease phenotypes of humans in response to SARS Cov-2 may provide more definitive information.

One limitation of our study is that we did not investigate the presence of virus in tissues of CVB3-infected mice. Similarly, it is unknown whether CFA administration can potentiate the production of protective neutralizing antibodies to CVB3, and if so, how long such an effect would last against different doses of virus. Likewise, whether administration of CFA in the face of CVB3 infection can mitigate the disease process is also unknown. At the time of this writing, we could not execute these experiments since our institutional guidelines do not allow any new animal experiments because of the ongoing threat of COVID-19 to the public. Nonetheless, our data may provide insights into our understanding of the occurrence of viral infections in the face of pre-existing, non-antigen-specific immune responses generated in response to a potentially wide range of environmental pathogens/microbes or gut microbiota over a period of time, which also may include formation of virtual memory cells [57, 58].

## Abbreviations

BCG: Bacillus Calmette-Guérin
CFA: complete Freund’s adjuvant
COVID-19: Coronavirus disease-19
CVB: Coxsackievirus B
EMEM: Eagle’s minimum essential medium
FBS: fetal bovine serum
IFN: interferon
I.p.: intra peritoneal
M.tb: *Mycobacterium tuberculosis*
NK: Natural killer
PBS: phosphate-buffered saline
RNA: ribonucleic acid
SARS-CoV-2: severe acute respiratory syndrome coronavirus 2
T1D Type: 1 diabetes
Th: T helper
VP: viral protein

## Author Contributions

Conceptualization, AG, CM and JR; validation, NL, DS, and JR; formal analysis, NL and DS; investigation, AG, CM, NL, DS and JR; data curation, NL; writing—original draft preparation, NL, AG, CM, and JR; writing—review and editing, NL, AG, CM, and JR; visualization, NL; funding acquisition, J.R.

## Acknowledgments

We thank Dr. Yuzhen Zhou for his assistance with the statistical analysis.

## Supplementary information

**Supplementary Table 1:**
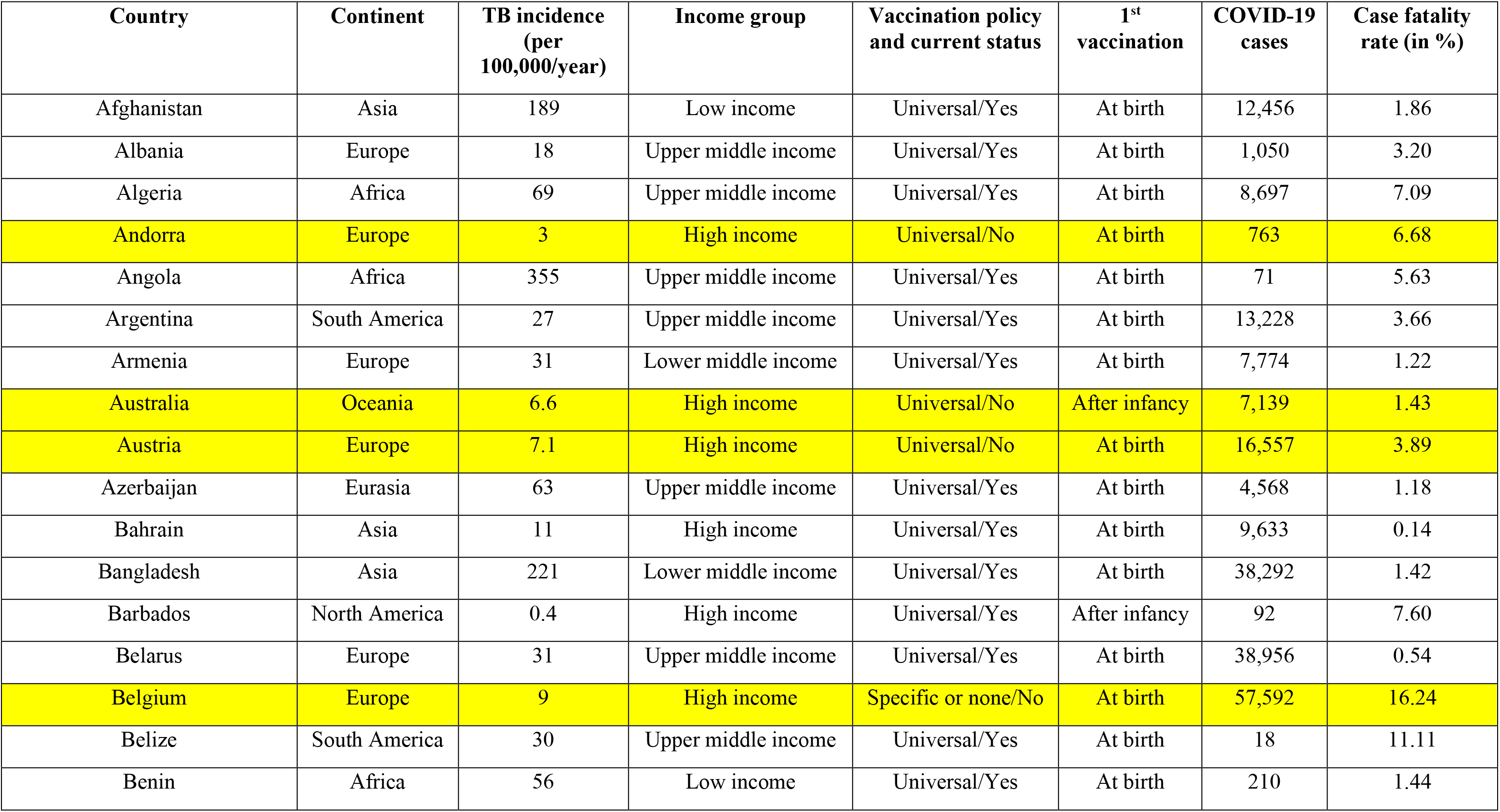
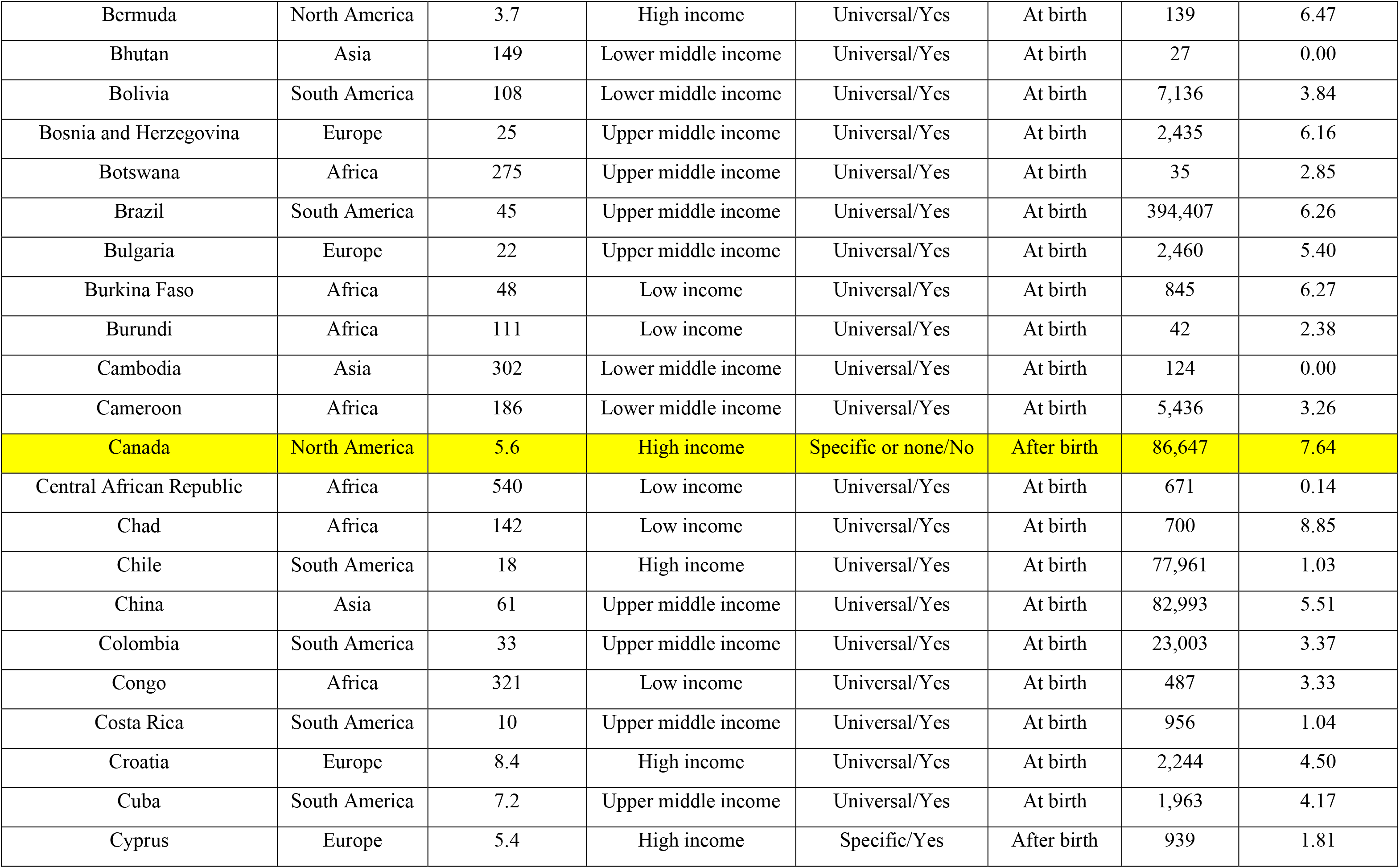
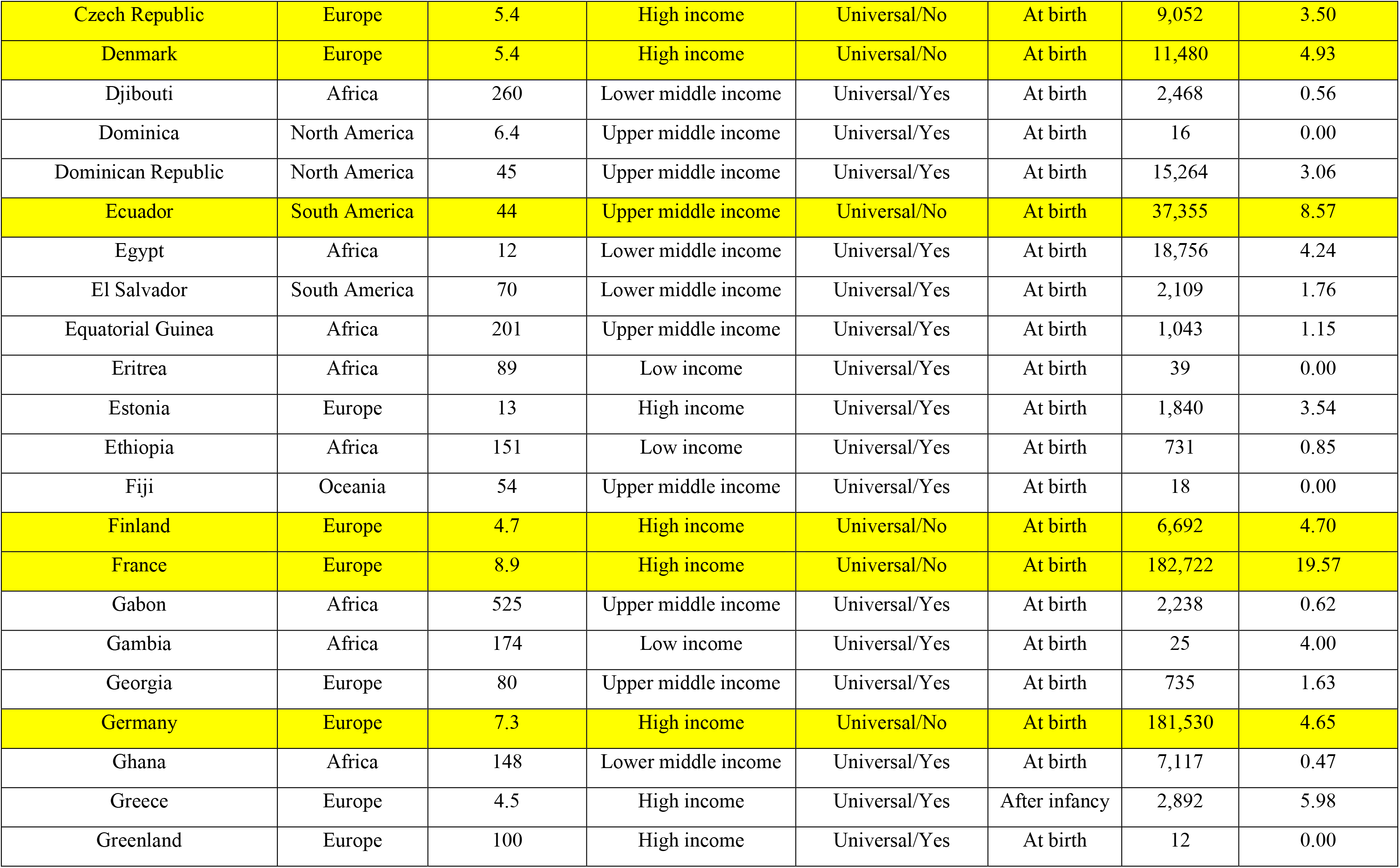
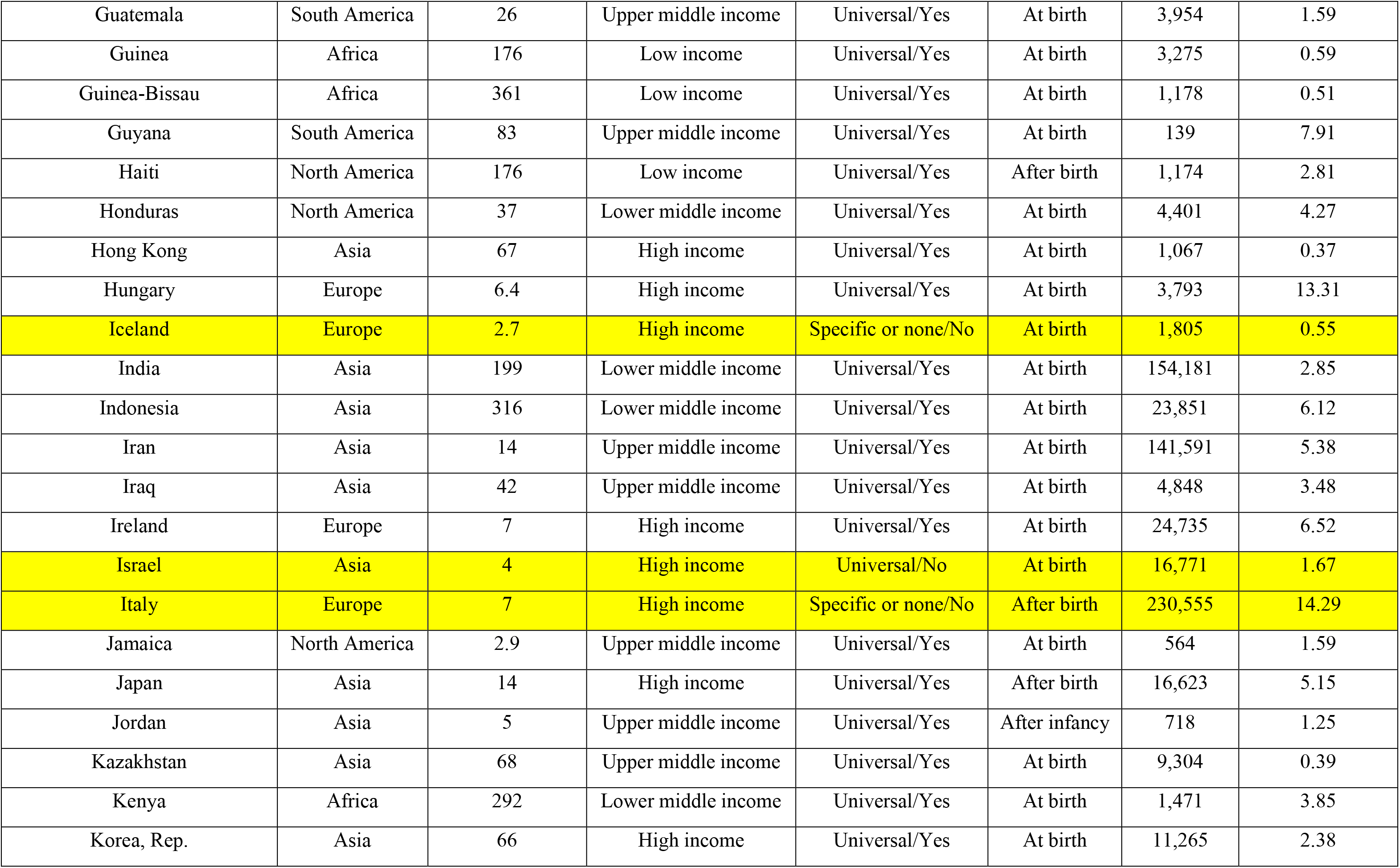
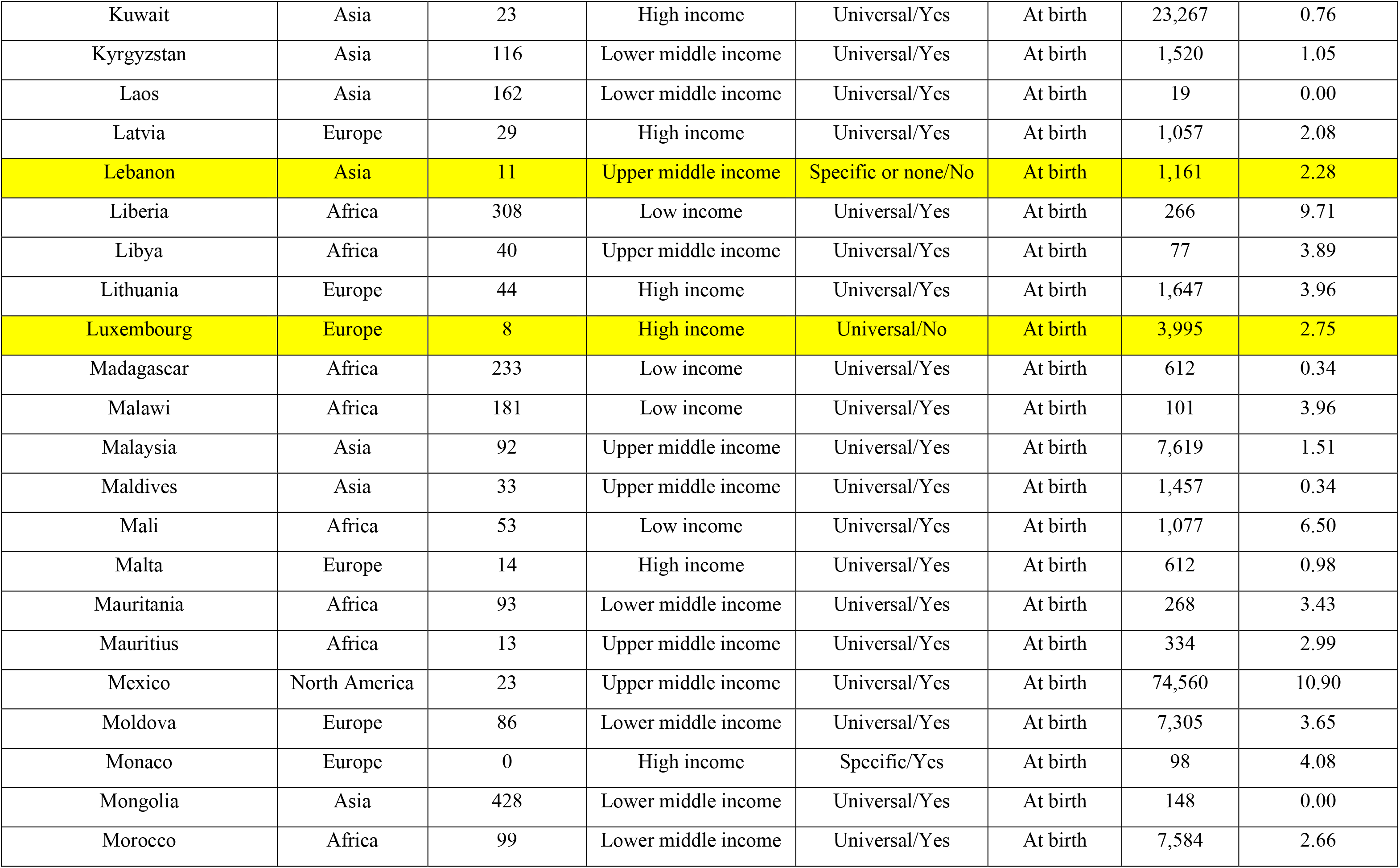
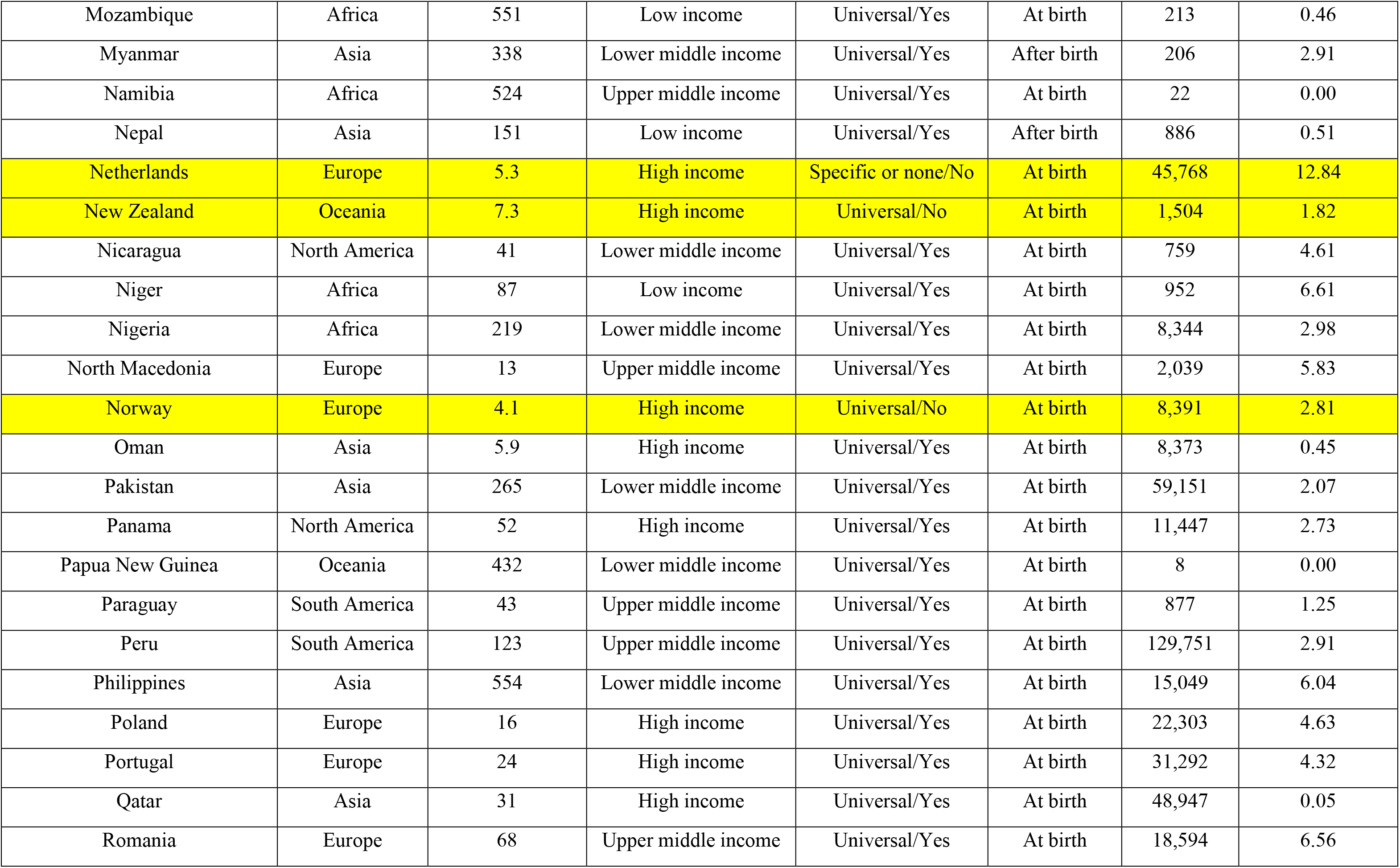
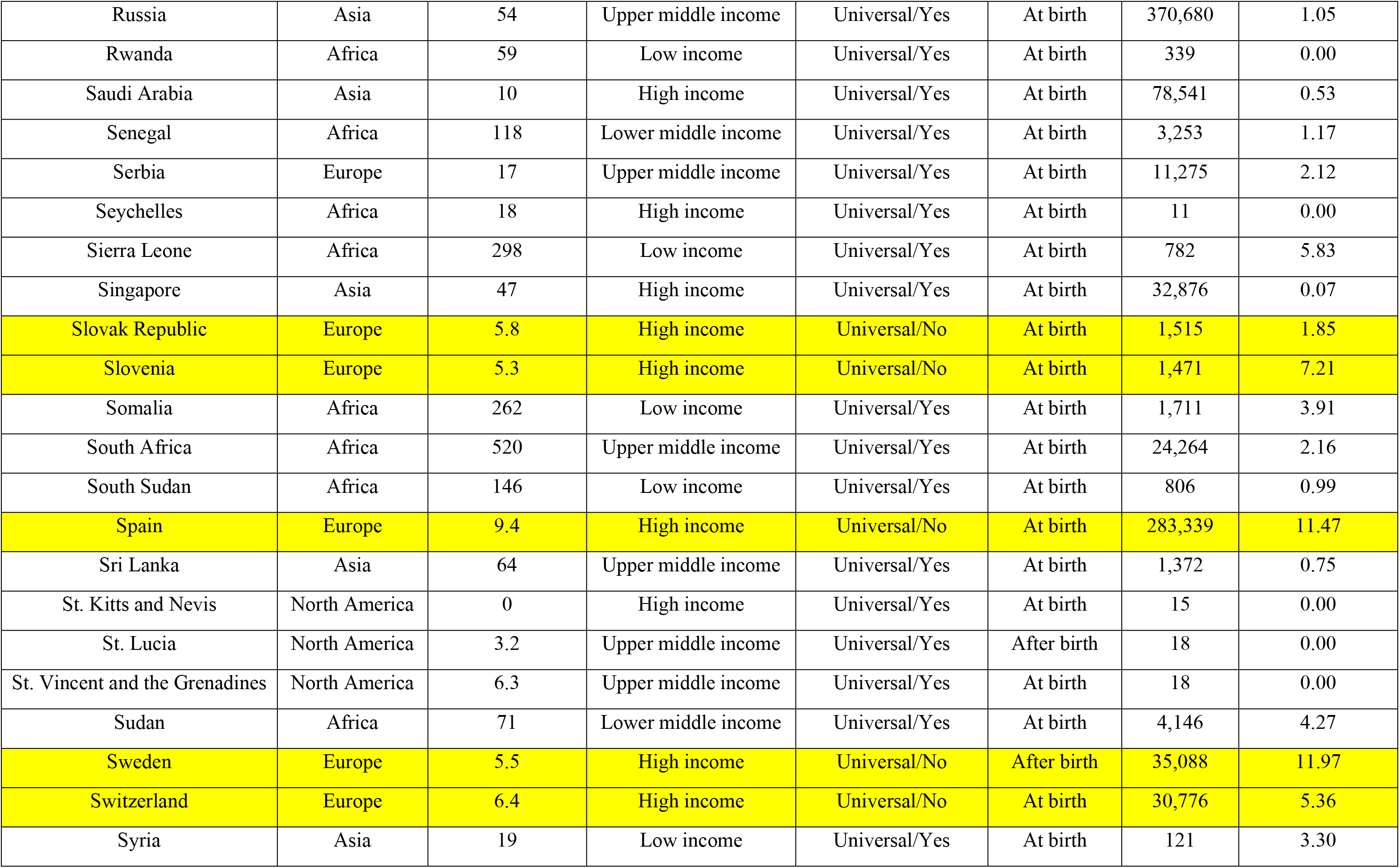
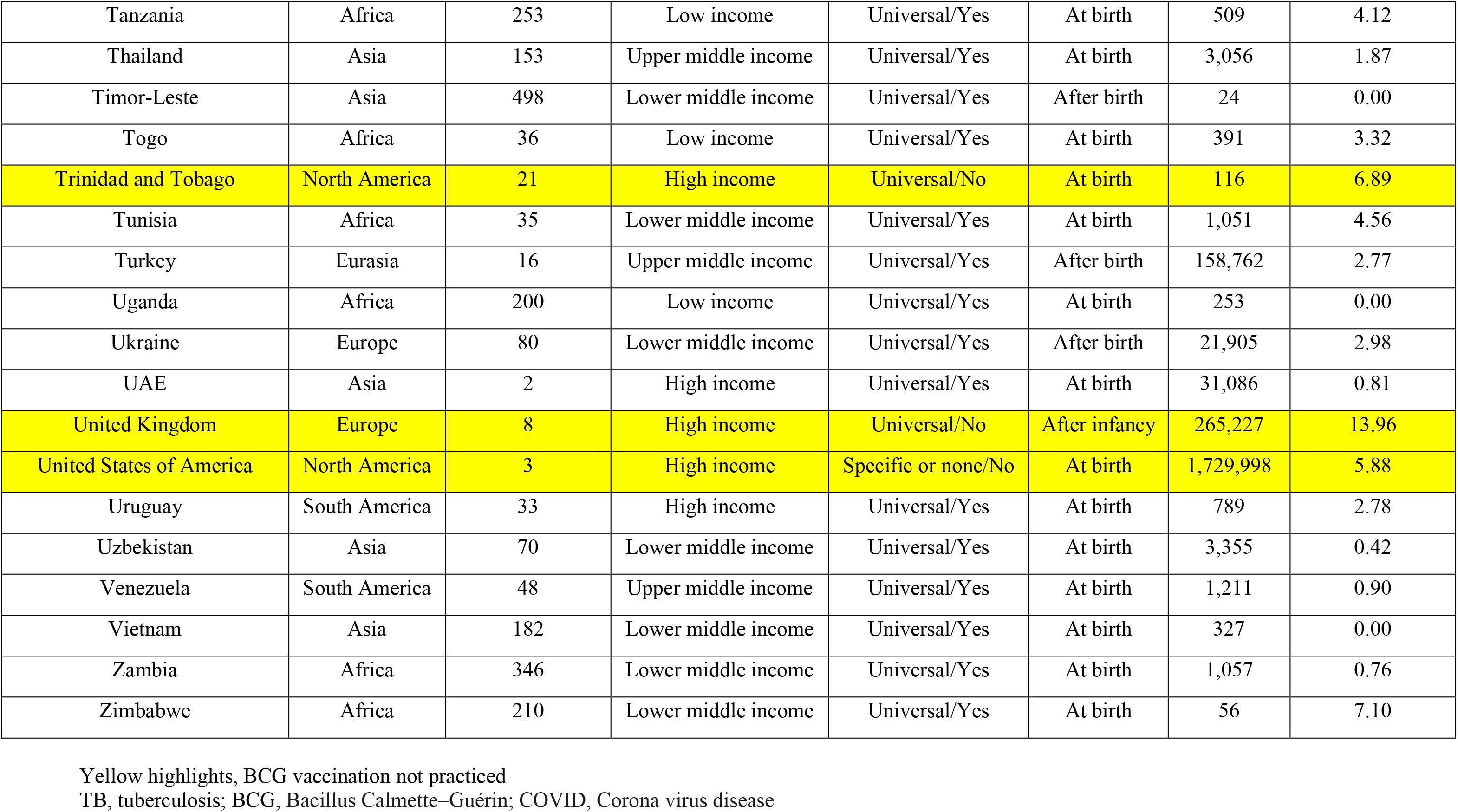
Current global status of BCG vaccination and its relationship with the number of COVID-19 cases reported as of May 27, 2020

