## Supplementary Table 1 for "Evidence for anti-viral effects of complete Freund’s adjuvant in the mouse model of enterovirus infection"

| **Supplementary Table 1: Current global status of BCG vaccination and its relationship with the number of COVID-19 cases reported as of May 27, 2020** | | | | | | | |
| --- | --- | --- | --- | --- | --- | --- | --- |
| **Country** | **Continent** | **TB incidence (per 100,000/year)** | **Income group** | **Vaccination policy and current status** | **1^st^ vaccination** | **COVID-19 cases** | **Case fatality rate (in %)** |
| Afghanistan | Asia | 189 | Low income | Universal/Yes | At birth | 12,456 | 1.86 |
| Albania | Europe | 18 | Upper middle income | Universal/Yes | At birth | 1,050 | 3.20 |
| Algeria | Africa | 69 | Upper middle income | Universal/Yes | At birth | 8,697 | 7.09 |
| Andorra | Europe | 3 | High income | Universal/No | At birth | 763 | 6.68 |
| Angola | Africa | 355 | Upper middle income | Universal/Yes | At birth | 71 | 5.63 |
| Argentina | South America | 27 | Upper middle income | Universal/Yes | At birth | 13,228 | 3.66 |
| Armenia | Europe | 31 | Lower middle income | Universal/Yes | At birth | 7,774 | 1.22 |
| Australia | Oceania | 6.6 | High income | Universal/No | After infancy | 7,139 | 1.43 |
| Austria | Europe | 7.1 | High income | Universal/No | At birth | 16,557 | 3.89 |
| Azerbaijan | Eurasia | 63 | Upper middle income | Universal/Yes | At birth | 4,568 | 1.18 |
| Bahrain | Asia | 11 | High income | Universal/Yes | At birth | 9,633 | 0.14 |
| Bangladesh | Asia | 221 | Lower middle income | Universal/Yes | At birth | 38,292 | 1.42 |
| Barbados | North America | 0.4 | High income | Universal/Yes | After infancy | 92 | 7.60 |
| Belarus | Europe | 31 | Upper middle income | Universal/Yes | At birth | 38,956 | 0.54 |
| Belgium | Europe | 9 | High income | Specific or none/No | At birth | 57,592 | 16.24 |
| Belize | South America | 30 | Upper middle income | Universal/Yes | At birth | 18 | 11.11 |
| Benin | Africa | 56 | Low income | Universal/Yes | At birth | 210 | 1.44 |
| Bermuda | North America | 3.7 | High income | Universal/Yes | At birth | 139 | 6.47 |
| Bhutan | Asia | 149 | Lower middle income | Universal/Yes | At birth | 27 | 0.00 |
| Bolivia | South America | 108 | Lower middle income | Universal/Yes | At birth | 7,136 | 3.84 |
| Bosnia and Herzegovina | Europe | 25 | Upper middle income | Universal/Yes | At birth | 2,435 | 6.16 |
| Botswana | Africa | 275 | Upper middle income | Universal/Yes | At birth | 35 | 2.85 |
| Brazil | South America | 45 | Upper middle income | Universal/Yes | At birth | 394,407 | 6.26 |
| Bulgaria | Europe | 22 | Upper middle income | Universal/Yes | At birth | 2,460 | 5.40 |
| Burkina Faso | Africa | 48 | Low income | Universal/Yes | At birth | 845 | 6.27 |
| Burundi | Africa | 111 | Low income | Universal/Yes | At birth | 42 | 2.38 |
| Cambodia | Asia | 302 | Lower middle income | Universal/Yes | At birth | 124 | 0.00 |
| Cameroon | Africa | 186 | Lower middle income | Universal/Yes | At birth | 5,436 | 3.26 |
| Canada | North America | 5.6 | High income | Specific or none/No | After birth | 86,647 | 7.64 |
| Central African Republic | Africa | 540 | Low income | Universal/Yes | At birth | 671 | 0.14 |
| Chad | Africa | 142 | Low income | Universal/Yes | At birth | 700 | 8.85 |
| Chile | South America | 18 | High income | Universal/Yes | At birth | 77,961 | 1.03 |
| China | Asia | 61 | Upper middle income | Universal/Yes | At birth | 82,993 | 5.51 |
| Colombia | South America | 33 | Upper middle income | Universal/Yes | At birth | 23,003 | 3.37 |
| Congo | Africa | 321 | Low income | Universal/Yes | At birth | 487 | 3.33 |
| Costa Rica | South America | 10 | Upper middle income | Universal/Yes | At birth | 956 | 1.04 |
| Croatia | Europe | 8.4 | High income | Universal/Yes | At birth | 2,244 | 4.50 |
| Cuba | South America | 7.2 | Upper middle income | Universal/Yes | At birth | 1,963 | 4.17 |
| Cyprus | Europe | 5.4 | High income | Specific/Yes | After birth | 939 | 1.81 |
| Czech Republic | Europe | 5.4 | High income | Universal/No | At birth | 9,052 | 3.50 |
| Denmark | Europe | 5.4 | High income | Universal/No | At birth | 11,480 | 4.93 |
| Djibouti | Africa | 260 | Lower middle income | Universal/Yes | At birth | 2,468 | 0.56 |
| Dominica | North America | 6.4 | Upper middle income | Universal/Yes | At birth | 16 | 0.00 |
| Dominican Republic | North America | 45 | Upper middle income | Universal/Yes | At birth | 15,264 | 3.06 |
| Ecuador | South America | 44 | Upper middle income | Universal/No | At birth | 37,355 | 8.57 |
| Egypt | Africa | 12 | Lower middle income | Universal/Yes | At birth | 18,756 | 4.24 |
| El Salvador | South America | 70 | Lower middle income | Universal/Yes | At birth | 2,109 | 1.76 |
| Equatorial Guinea | Africa | 201 | Upper middle income | Universal/Yes | At birth | 1,043 | 1.15 |
| Eritrea | Africa | 89 | Low income | Universal/Yes | At birth | 39 | 0.00 |
| Estonia | Europe | 13 | High income | Universal/Yes | At birth | 1,840 | 3.54 |
| Ethiopia | Africa | 151 | Low income | Universal/Yes | At birth | 731 | 0.85 |
| Fiji | Oceania | 54 | Upper middle income | Universal/Yes | At birth | 18 | 0.00 |
| Finland | Europe | 4.7 | High income | Universal/No | At birth | 6,692 | 4.70 |
| France | Europe | 8.9 | High income | Universal/No | At birth | 182,722 | 19.57 |
| Gabon | Africa | 525 | Upper middle income | Universal/Yes | At birth | 2,238 | 0.62 |
| Gambia | Africa | 174 | Low income | Universal/Yes | At birth | 25 | 4.00 |
| Georgia | Europe | 80 | Upper middle income | Universal/Yes | At birth | 735 | 1.63 |
| Germany | Europe | 7.3 | High income | Universal/No | At birth | 181,530 | 4.65 |
| Ghana | Africa | 148 | Lower middle income | Universal/Yes | At birth | 7,117 | 0.47 |
| Greece | Europe | 4.5 | High income | Universal/Yes | After infancy | 2,892 | 5.98 |
| Greenland | Europe | 100 | High income | Universal/Yes | At birth | 12 | 0.00 |
| Guatemala | South America | 26 | Upper middle income | Universal/Yes | At birth | 3,954 | 1.59 |
| Guinea | Africa | 176 | Low income | Universal/Yes | At birth | 3,275 | 0.59 |
| Guinea-Bissau | Africa | 361 | Low income | Universal/Yes | At birth | 1,178 | 0.51 |
| Guyana | South America | 83 | Upper middle income | Universal/Yes | At birth | 139 | 7.91 |
| Haiti | North America | 176 | Low income | Universal/Yes | After birth | 1,174 | 2.81 |
| Honduras | North America | 37 | Lower middle income | Universal/Yes | At birth | 4,401 | 4.27 |
| Hong Kong | Asia | 67 | High income | Universal/Yes | At birth | 1,067 | 0.37 |
| Hungary | Europe | 6.4 | High income | Universal/Yes | At birth | 3,793 | 13.31 |
| Iceland | Europe | 2.7 | High income | Specific or none/No | At birth | 1,805 | 0.55 |
| India | Asia | 199 | Lower middle income | Universal/Yes | At birth | 154,181 | 2.85 |
| Indonesia | Asia | 316 | Lower middle income | Universal/Yes | At birth | 23,851 | 6.12 |
| Iran | Asia | 14 | Upper middle income | Universal/Yes | At birth | 141,591 | 5.38 |
| Iraq | Asia | 42 | Upper middle income | Universal/Yes | At birth | 4,848 | 3.48 |
| Ireland | Europe | 7 | High income | Universal/Yes | At birth | 24,735 | 6.52 |
| Israel | Asia | 4 | High income | Universal/No | At birth | 16,771 | 1.67 |
| Italy | Europe | 7 | High income | Specific or none/No | After birth | 230,555 | 14.29 |
| Jamaica | North America | 2.9 | Upper middle income | Universal/Yes | At birth | 564 | 1.59 |
| Japan | Asia | 14 | High income | Universal/Yes | After birth | 16,623 | 5.15 |
| Jordan | Asia | 5 | Upper middle income | Universal/Yes | After infancy | 718 | 1.25 |
| Kazakhstan | Asia | 68 | Upper middle income | Universal/Yes | At birth | 9,304 | 0.39 |
| Kenya | Africa | 292 | Lower middle income | Universal/Yes | At birth | 1,471 | 3.85 |
| Korea, Rep. | Asia | 66 | High income | Universal/Yes | At birth | 11,265 | 2.38 |
| Kuwait | Asia | 23 | High income | Universal/Yes | At birth | 23,267 | 0.76 |
| Kyrgyzstan | Asia | 116 | Lower middle income | Universal/Yes | At birth | 1,520 | 1.05 |
| Laos | Asia | 162 | Lower middle income | Universal/Yes | At birth | 19 | 0.00 |
| Latvia | Europe | 29 | High income | Universal/Yes | At birth | 1,057 | 2.08 |
| Lebanon | Asia | 11 | Upper middle income | Specific or none/No | At birth | 1,161 | 2.28 |
| Liberia | Africa | 308 | Low income | Universal/Yes | At birth | 266 | 9.71 |
| Libya | Africa | 40 | Upper middle income | Universal/Yes | At birth | 77 | 3.89 |
| Lithuania | Europe | 44 | High income | Universal/Yes | At birth | 1,647 | 3.96 |
| Luxembourg | Europe | 8 | High income | Universal/No | At birth | 3,995 | 2.75 |
| Madagascar | Africa | 233 | Low income | Universal/Yes | At birth | 612 | 0.34 |
| Malawi | Africa | 181 | Low income | Universal/Yes | At birth | 101 | 3.96 |
| Malaysia | Asia | 92 | Upper middle income | Universal/Yes | At birth | 7,619 | 1.51 |
| Maldives | Asia | 33 | Upper middle income | Universal/Yes | At birth | 1,457 | 0.34 |
| Mali | Africa | 53 | Low income | Universal/Yes | At birth | 1,077 | 6.50 |
| Malta | Europe | 14 | High income | Universal/Yes | At birth | 612 | 0.98 |
| Mauritania | Africa | 93 | Lower middle income | Universal/Yes | At birth | 268 | 3.43 |
| Mauritius | Africa | 13 | Upper middle income | Universal/Yes | At birth | 334 | 2.99 |
| Mexico | North America | 23 | Upper middle income | Universal/Yes | At birth | 74,560 | 10.90 |
| Moldova | Europe | 86 | Lower middle income | Universal/Yes | At birth | 7,305 | 3.65 |
| Monaco | Europe | 0 | High income | Specific/Yes | At birth | 98 | 4.08 |
| Mongolia | Asia | 428 | Lower middle income | Universal/Yes | At birth | 148 | 0.00 |
| Morocco | Africa | 99 | Lower middle income | Universal/Yes | At birth | 7,584 | 2.66 |
| Mozambique | Africa | 551 | Low income | Universal/Yes | At birth | 213 | 0.46 |
| Myanmar | Asia | 338 | Lower middle income | Universal/Yes | After birth | 206 | 2.91 |
| Namibia | Africa | 524 | Upper middle income | Universal/Yes | At birth | 22 | 0.00 |
| Nepal | Asia | 151 | Low income | Universal/Yes | After birth | 886 | 0.51 |
| Netherlands | Europe | 5.3 | High income | Specific or none/No | At birth | 45,768 | 12.84 |
| New Zealand | Oceania | 7.3 | High income | Universal/No | At birth | 1,504 | 1.82 |
| Nicaragua | North America | 41 | Lower middle income | Universal/Yes | At birth | 759 | 4.61 |
| Niger | Africa | 87 | Low income | Universal/Yes | At birth | 952 | 6.61 |
| Nigeria | Africa | 219 | Lower middle income | Universal/Yes | At birth | 8,344 | 2.98 |
| North Macedonia | Europe | 13 | Upper middle income | Universal/Yes | At birth | 2,039 | 5.83 |
| Norway | Europe | 4.1 | High income | Universal/No | At birth | 8,391 | 2.81 |
| Oman | Asia | 5.9 | High income | Universal/Yes | At birth | 8,373 | 0.45 |
| Pakistan | Asia | 265 | Lower middle income | Universal/Yes | At birth | 59,151 | 2.07 |
| Panama | North America | 52 | High income | Universal/Yes | At birth | 11,447 | 2.73 |
| Papua New Guinea | Oceania | 432 | Lower middle income | Universal/Yes | At birth | 8 | 0.00 |
| Paraguay | South America | 43 | Upper middle income | Universal/Yes | At birth | 877 | 1.25 |
| Peru | South America | 123 | Upper middle income | Universal/Yes | At birth | 129,751 | 2.91 |
| Philippines | Asia | 554 | Lower middle income | Universal/Yes | At birth | 15,049 | 6.04 |
| Poland | Europe | 16 | High income | Universal/Yes | At birth | 22,303 | 4.63 |
| Portugal | Europe | 24 | High income | Universal/Yes | At birth | 31,292 | 4.32 |
| Qatar | Asia | 31 | High income | Universal/Yes | At birth | 48,947 | 0.05 |
| Romania | Europe | 68 | Upper middle income | Universal/Yes | At birth | 18,594 | 6.56 |
| Russia | Asia | 54 | Upper middle income | Universal/Yes | At birth | 370,680 | 1.05 |
| Rwanda | Africa | 59 | Low income | Universal/Yes | At birth | 339 | 0.00 |
| Saudi Arabia | Asia | 10 | High income | Universal/Yes | At birth | 78,541 | 0.53 |
| Senegal | Africa | 118 | Lower middle income | Universal/Yes | At birth | 3,253 | 1.17 |
| Serbia | Europe | 17 | Upper middle income | Universal/Yes | At birth | 11,275 | 2.12 |
| Seychelles | Africa | 18 | High income | Universal/Yes | At birth | 11 | 0.00 |
| Sierra Leone | Africa | 298 | Low income | Universal/Yes | At birth | 782 | 5.83 |
| Singapore | Asia | 47 | High income | Universal/Yes | At birth | 32,876 | 0.07 |
| Slovak Republic | Europe | 5.8 | High income | Universal/No | At birth | 1,515 | 1.85 |
| Slovenia | Europe | 5.3 | High income | Universal/No | At birth | 1,471 | 7.21 |
| Somalia | Africa | 262 | Low income | Universal/Yes | At birth | 1,711 | 3.91 |
| South Africa | Africa | 520 | Upper middle income | Universal/Yes | At birth | 24,264 | 2.16 |
| South Sudan | Africa | 146 | Low income | Universal/Yes | At birth | 806 | 0.99 |
| Spain | Europe | 9.4 | High income | Universal/No | At birth | 283,339 | 11.47 |
| Sri Lanka | Asia | 64 | Upper middle income | Universal/Yes | At birth | 1,372 | 0.75 |
| St. Kitts and Nevis | North America | 0 | High income | Universal/Yes | At birth | 15 | 0.00 |
| St. Lucia | North America | 3.2 | Upper middle income | Universal/Yes | After birth | 18 | 0.00 |
| St. Vincent and the Grenadines | North America | 6.3 | Upper middle income | Universal/Yes | At birth | 18 | 0.00 |
| Sudan | Africa | 71 | Lower middle income | Universal/Yes | At birth | 4,146 | 4.27 |
| Sweden | Europe | 5.5 | High income | Universal/No | After birth | 35,088 | 11.97 |
| Switzerland | Europe | 6.4 | High income | Universal/No | At birth | 30,776 | 5.36 |
| Syria | Asia | 19 | Low income | Universal/Yes | At birth | 121 | 3.30 |
| Tanzania | Africa | 253 | Low income | Universal/Yes | At birth | 509 | 4.12 |
| Thailand | Asia | 153 | Upper middle income | Universal/Yes | At birth | 3,056 | 1.87 |
| Timor-Leste | Asia | 498 | Lower middle income | Universal/Yes | After birth | 24 | 0.00 |
| Togo | Africa | 36 | Low income | Universal/Yes | At birth | 391 | 3.32 |
| Trinidad and Tobago | North America | 21 | High income | Universal/No | At birth | 116 | 6.89 |
| Tunisia | Africa | 35 | Lower middle income | Universal/Yes | At birth | 1,051 | 4.56 |
| Turkey | Eurasia | 16 | Upper middle income | Universal/Yes | After birth | 158,762 | 2.77 |
| Uganda | Africa | 200 | Low income | Universal/Yes | At birth | 253 | 0.00 |
| Ukraine | Europe | 80 | Lower middle income | Universal/Yes | After birth | 21,905 | 2.98 |
| UAE | Asia | 2 | High income | Universal/Yes | At birth | 31,086 | 0.81 |
| United Kingdom | Europe | 8 | High income | Universal/No | After infancy | 265,227 | 13.96 |
| United States of America | North America | 3 | High income | Specific or none/No | At birth | 1,729,998 | 5.88 |
| Uruguay | South America | 33 | High income | Universal/Yes | At birth | 789 | 2.78 |
| Uzbekistan | Asia | 70 | Lower middle income | Universal/Yes | At birth | 3,355 | 0.42 |
| Venezuela | South America | 48 | Upper middle income | Universal/Yes | At birth | 1,211 | 0.90 |
| Vietnam | Asia | 182 | Lower middle income | Universal/Yes | At birth | 327 | 0.00 |
| Zambia | Africa | 346 | Lower middle income | Universal/Yes | At birth | 1,057 | 0.76 |
| Zimbabwe | Africa | 210 | Lower middle income | Universal/Yes | At birth | 56 | 7.10 |

Yellow highlights, BCG vaccination not practiced

TB, tuberculosis; BCG, Bacillus Calmette–Guérin; COVID, Corona virus disease
